# Force transmission and SUN-KASH higher-order assembly in the LINC complex models

**DOI:** 10.1101/2023.02.02.526920

**Authors:** Ghafar Yerima, Nya Domkam, Jessica Ornowski, Zeinab Jahed, Mohammad R.K. Mofrad

**Author notes:** These authors contributed equally to this work (Ghafar Yerima, Nya Domkam).

## Abstract

The linker of the nucleoskeleton and cytoskeleton (LINC) complex comprises SUN (Sad-1 and UNC-84) and KASH (Klarsicht, ANC-1, SYNE homology) domain proteins, whose conserved interactions provide a physical coupling between the cytoskeleton and the nucleoskeleton, thereby mediating the transfer of physical forces across the nuclear envelope. The LINC complex can perform distinct cellular functions by pairing various KASH domain proteins with the same SUN domain protein. Recent studies have suggested a higher-order assembly of SUN and KASH instead of a more widely accepted linear trimer model for the LINC complex. In the present study, we use molecular dynamics simulations to investigate the mechanism of force transfer across the two proposed models of LINC complex assembly, namely the 3:3 linear trimer model and the 6:6 higher-order model. Employing steered molecular dynamics simulations with various structures using forces at different rates and directions, we examine the structural stability of the two models under various biologically relevant conditions. Our results suggest that both models can withstand and transfer significant levels of force while retaining their structural integrity. However, the force response of various SUN KASH assemblies depended on the force direction and pulling rates. Slower pulling rates resulted in higher mean square fluctuations of the 3:3 assembly compared to the fast pulling. Interestingly, the 6:6 assembly tends to provide an additional range of motion flexibility and might be more suitable for describing the interaction between SUN and KASH under compressive and shear forces. These findings offer insights into how the SUN and KASH proteins maintain the structural integrity of the nuclear membrane.

## Introduction

The LINC (**LI**nkers of **N**ucleoskeleton and **C**ytoskeleton) complex is a vast network of proteins involved in the mechanical response of the cell ^1–3^. The main components of the LINC complex are the SUN (**S**ad1 and **UN**C-84) and KASH (**K**larsicht, **A**NC-1 and **S**YNE/Nesprin-1 and -2 **H**omology) domain proteins ^4^. SUN proteins are anchored to the inner nuclear membrane and contain small domains in the nucleoplasm and large domains that extend to the outer nuclear membrane. KASH proteins contain a small KASH domain in the perinuclear space and large domains that extend into the cytoplasm ^5,6^. Both SUN and KASH proteins can transduce forces from the cytoplasm to the nucleoplasm ^7–9^. Of the five different SUN proteins, only two, namely SUN1 and SUN2, are commonly found in virtually all cells^10^. KASH domain proteins function not only inside the nuclear envelope, but also in the cytoplasm. There are at least six mammalian KASH domain proteins, namely Nesprins 1-4, KASH 5, and lymphoid-restricted membrane proteins (^11,12,13,14^). Nesprin 1 and 2 are widely expressed in most cell types, whereas Nesprin 3, 4, & KASH5 are cell type-specific. KASH can interact with various cytoskeletal elements: F-actin ^15^, microtubules ^16^, intermediate filaments via a plectin binding site ^17^, and the dynein-dynactin ^18^ complexes which perform specific cell functions.

Mutations in SUN and KASH proteins are believed to lead to a variety of structural and functional defects in the cell and have been linked to various human diseases ^19–23^. For instance, Nesprins mutations are associated with autosomal recessive cerebellar ataxia^24^, recessive arthrogryposis multiplex congenita ^25^, hearing loss ^26^, Meckel–Gruber syndrome ^27^, depression, and bipolar disorder ^28^. SUN mutations are associated with DYT1 dystonia ^29^ and muscular dystrophies including Emery-Dreifuss muscular dystrophy (EDMD)^30^.

Several studies by our group and others have aimed at understanding the molecular mechanisms of force transfer across SUN/KASH complexes through experimental and computational techniques. Notable advancements in our understanding of the LINC complex molecular structure were made by the solved crystal structure of the conserved SUN2/KASH2 interaction by three independent groups a decade ago ^5,31,32^. The crystal structure revealed an arrangement referred to as the 3:3 model meaning that each SUN forms a trimer to interact with 3 KASH, forming an overall hexamer (**Figure 1A, B**). These structural findings suggested that SUN/KASH arrange as linear arrays in the NE ^31^. Studies based on these crystal structures revealed several details regarding the molecular mechanisms of force transfer across the SUN/KASH complex. For instance, recent studies using combined *in silico* molecular dynamics simulations and *in vivo C*.*elegans genetics* showed that a mutation of tyrosine at position -7 of KASH disrupts SUN/KASH interaction ^33^. We showed that a disulfide bond is required for maximal force transmission in SUN2/KASH1 & 2 ^34^, and different KASH proteins may transfer distinct magnitudes of force ^35^. However, all the above-mentioned studies were based on the linear trimer model of SUN/KASH, using the SUN2/KASH1 & 2 structures. Recently, new crystal structures of the SUN/KASH interaction were released by two groups ^36,37^. These new structures suggest an alternative model in which SUN/KASH pairs may form higher-order assemblies instead of the putative linear trimer model^36^. This alternative arrangement is known as the 6:6, in which two SUN/KASH hexamers (3:3 arrangements) interact through their SUN and KASH domains in a head-to-head fashion and form an overall dodecamer (**Figure 1A, C**).

**Figure 1:**
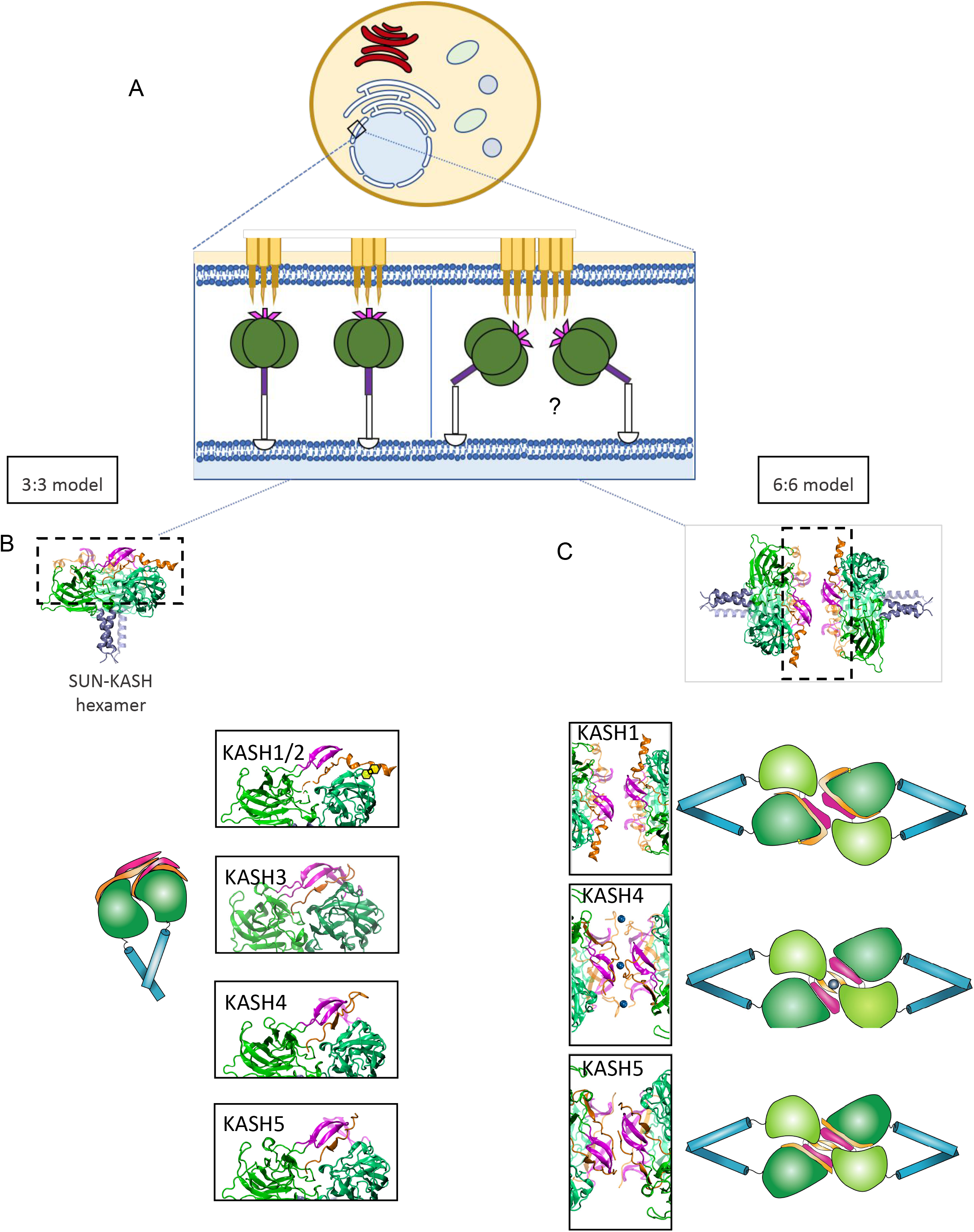
Linear trimer model and high-order assembly of SUN1,2 in complex with various KASH. **A)** Schematic of a cell with an enlarged view of the nuclear membrane which displays the main components of the LINC complex. From top to bottom of the enlarged view, we have the Nesprins and KASH domains (orange), the KASH-lids (pink), the SUN domains (green), and the coiled-coils (purple). **B**) Crystal structure of the linear trimer model or 3:3 model with the different SUN2/KASH pairs. The right column shows the adjacent SUN domains (green), the KASH domain (orange) as well as the KASH-lids (pink) of KASH1,2, KASH3, KASH4, and KASH5. A disulfide bond between the SUN domain and the neighboring KASH is represented in yellow for KASH1,2. **C**) Crystal structure of the high-order assembly or 6:6 model with the different SUN1/KASH pairs and an enlarged view of the head-to-head interaction. The right column shows the SUN domains (green), the KASH domain (orange) as well as the KASH-lids (pink) of KASH1, KASH4, and KASH5.

The 6:6 assembly was solved for three SUN/KASH pairs, namely SUN1/KASH1, SUN1/KASH4, and SUN1/KASH5^36^. Each one of these complexes involve a unique binding modality. In the SUN1/KASH1 complex, the KASH-lid, a region within a SUN protomer, maintains the 6:6 head-to-head interaction. Zinc-cysteine coordination bonds in SUN1/ maintain the head-to-head interaction. SUN1/KASH5, like SUN1/KASH1, employs the KASH-lid to preserve its head-to-head interaction. The only difference is an additional interaction between the KASH domains of the opposing SUN/KASH hexamers, which is mediated by a PPP motif ^36^. Additionally, novel 3:3 complexes were crystallized, namely SUN2/KASH3, SUN2/KASH4, and SUN2/KASH5 configurations ^37^. Unlike KASH 1 & 2 which form a 90° angle due to a proline on the -11th position, the other 3 KASH do not make a 90° kink and lie along the SUN protomer (**Figure 2A, B**).

**Figure 2:**
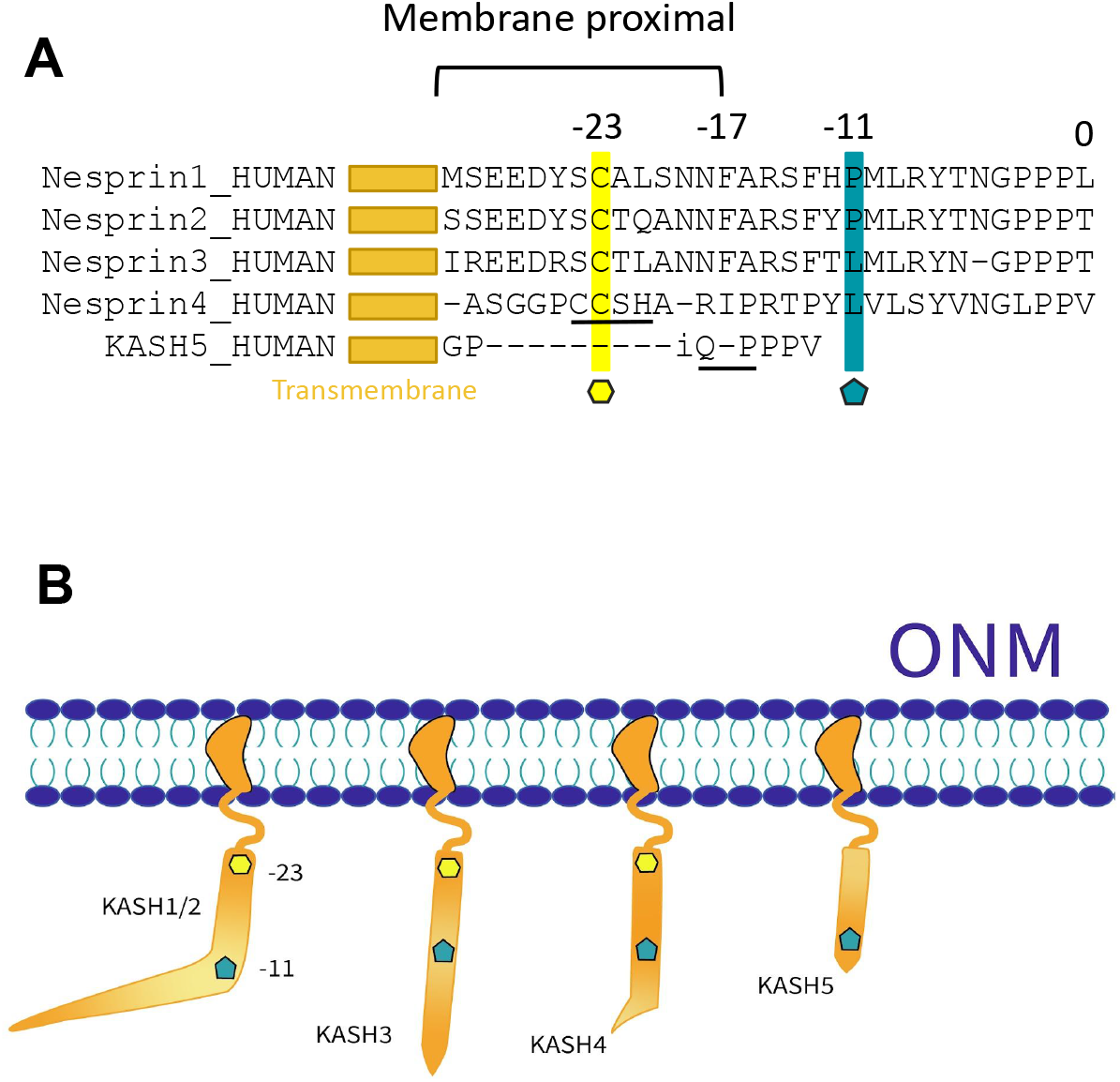
**A**) Multiple sequence alignment of various human KASH domain proteins. Position 0 marks the C terminus of the KASH domains. Position -11 corresponds to a proline in KASH1,2 and a leucine in KASH3,4,5 (blue pentagon). Position -23 is occupied by a cysteine residue in KASH1-4 (yellow hexagon). CCSH and PPP domains are found in KASH4 and KASH5 respectively (underlined). **B)** Schematic of the perinuclear space containing KASH domains. A 90-degree kink at proline -11 is observed in KASH1,2. This kink is not observed in the KASH3,4,5 structures.

Because force transfer across the LINC complex depends highly on how various LINC complexes assemble in the NE, a detailed evaluation and analysis of the two models is needed. To our knowledge, no studies have compared the molecular mechanisms of force transfer across the two models for various SUN/KASH pairs. Moreover, little is known about the response of the SUN/KASH complex to mechanical forces at varying directions and rates, conceivable at various cellular processes.

In this study, we develop all-atom molecular dynamics simulations to analyze and compare the mechanism(s) of force transfer across the two proposed models of LINC complex assembly, namely the 3:3 linear trimer model, and the 6:6 higher-order model. Specifically, we investigate SUN2 in complex with KASH1-5 and SUN1 in complex with KASH1, 4, & 5. Because of the similarity between SUN1 and SUN2, the structures in both models can be effectively compared based on the KASH type. Thus, we conduct molecular simulations to expose the structures to mechanical forces at different rates and directions and evaluate the structural stability of the two models under various biologically relevant conditions. We used uniaxial pulling for the 3:3 linear trimer and 6:6 higher-order model, and transverse pulling for the 6:6 higher-order model. By comparing the force response and conformational changes in each model, our results suggest that both models can withstand high forces without significant deformation. Moreover, the higher-order assembly provides an additional range of motion and might be more suitable for describing the interaction between SUN and KASH under compressive and shear forces. Ultimately, these findings offer insight into how the SUN and KASH proteins assemble in the nuclear membrane and maintain the structural integrity of the nuclear envelope.

## Materials and Methods

### Models of SUN/KASH complexes

Seven 3:3 and four 6:6 structures were downloaded from the Protein Data Bank and used for our simulations. The 3:3 structures were: SUN1/KASH4 (pdb code: **6R16**^36^), SUN1/KASH5 (pdb code: **6R2I**^36^), SUN2/KASH1 (pdb code: **4DXR**^31^), SUN2/KASH2 (pdb code: **4DXS**^31^), SUN2/KASH3 (pdb code: **6WME**^37^), SUN2/KASH4 (pdb code: **6WMD**^37^), and SUN2/KASH5 (pdb code: **6WMF**^37^). The SUN1 structures, which were originally 6:6 structures, were split in half to obtain their 3:3 versions using the Visual Molecular Dynamics (VMD) software. The four 6:6 structures were SUN1/KASH1 (pdb code: **6R15**^36^), SUN1/KASH4 (pdb code: **6R16**), SUN1/KASH5 (pdb code: **6R2I)**, and Apo-SUN2 (pdb code: **6WMG**^37^). The SUN1 and SUN2 protomers in each structure had similar lengths (195-196 aa) with a few missing residues. However, these residues were not near the regions of interest.

### Simulation Protocol

All simulations were performed using GROMACS ^38^ free software with the CHARMM36 ^39^ force field. All structures were minimized at 5000 steps with an energy tolerance of 1000 KJ/mol/nm and equilibrated for roughly 5 ns with a time step of 2 fs. These simulations were run at a constant temperature of 310 K with Berendsen temperature and pressure coupling. Periodic boundary conditions were applied in all three directions. Two different types of pulling simulations were performed on the structures using the Isothermal–isobaric ensemble. In total, 6 different pulling modalities were conducted involving specific pulling rates, pulling direction, and pulling groups. The structures were pulled at a constant velocity of 10 nm/ns for 0.5 ns and another set of simulations were pulled at 1 nm/ns for 5 ns to achieve the same displacement of 5 nm. For all 3:3 structures, the last residues on the different KASH proteins were pulled in the opposite direction of the CC region while the end residues of the SUN domains were fixed in all three dimensions. In the 6:6 structure, the nitrogen on one set of coiled regions was pulled in the opposite direction of the CC region. The KASH in the 6:6 structure in a different simulation series was pulled orthogonally to the direction of the coiled-coil domain (**Table1**).

**Table 1:** Simulation breakdown between the different structures in the 3:3 linear model and 6:6 higher-order assembly. *Only the uniaxial on KASH end residue modality were used for Apo-SUN2

|  | Structures | Pulling modality | Pulling Rate | Simulations run |
| --- | --- | --- | --- | --- |
| 3:3 Linear model | SUN1/KASH4 | Uniaxial on KASH end residue | 1 nm/ns for 5 ns | 3 |
|  | SUN1/KASH5 |  | + |  |
|  | SUN2/KASH1 |  |  |  |
|  | SUN2/KASH2 |  |  |  |
|  | SUN2/KASH3 |  | 10 nm/ns for 0.5 ns |  |
|  | SUN2/KASH4 |  |  |  |
|  | SUN2/KASH5 |  |  |  |
| 6:6 Higher-order assembly | SUN1/KASH1 | Transverse on KASH end residue | 1 nm/ns for 5 ns | 3 |
|  | SUN1/KASH4 |  | + |  |
|  | SUN1/KASH5 |  |  |  |
|  | Apo-SUN2* |  | 10 nm/ns for 0.5 ns |  |
|  |  | Uniaxial on KASH end residue |  |  |

### Post-Processing/Trajectory analyses

#### RMSD

Root mean square deviation of the protein backbone was calculated using GROMACS ^38^ free software and plotted using Xmgrace ^40^. Each frame in the trajectory was aligned and referenced with the first frame.

#### RMSF

Root mean square fluctuation per residue was calculated using GROMACS^38^ free software after fitting to the first frame of the pulling simulation and plotted using Python^41^. For the 3:3 model structures and for each pulling rate, the data for all SUN and KASH protomers of 3 simulations were averaged.

#### Pulling Force Quantification

The force graphs were obtained from the input files of each simulation using GROMACS^38^ free software and plotted using Python ^41^. The data for 3 simulations for each pulling rate were averaged per structure.

#### Interaction Energies

The short range nonbonded interaction energies for the salt bridge residue pairs (K533-E672, D542-R708) were calculated using GROMACS^38^ free software and plotted using Python ^41^. The salt bridges were categorized as intramolecular (D542-R708) and intermolecular (K533-E672). The data were then concatenated per salt bridge type for each simulation. The data for three simulations was concatenated and density plots were obtained.

#### Heatmap

Piecewise distances between residues were calculated between opposing residues on KASH-lids using VMD. The heatmap was created using the positional data and Python ^41^ library Seaborn.

#### Angle calculation

New conformational angle changes were obtained using VMD’s position tracker over the simulation time and Excel. The alpha carbons of residues 631 and 619 were used to create a vector, and all vectors on one head of SUN/KASH were averaged into one vector. The vector calculations were done through Excel.

#### Visualization

All visualizations were done using the Visual Molecular Dynamics (VMD) software ^42^.

## Results

Recent structural biological studies have proposed a higher-order assembly model for the LINC complex, casting doubts on the putative mechanism of force transfer within the SUN/KASH models. In this study, we used molecular dynamics simulations contrasting the putative 3:3 SUN/KASH complex model and the new higher-order assembly 6:6 SUN/KASH complex. Both models were subjected to pulling forces at different pulling rates and directions to mimic biologically relevant conditions. These results suggest the assembly of a more profound SUN/ KASH network in the nuclear envelope.

### Mechanics of Force Transfer in the 3:3 SUN/KASH Complex

The 3:3 SUN/KASH complex was exposed to pulling forces in the uniaxial direction at two rates: 1 nm/ns for 5 ns and 10 nm/ns for 0.5 ns. The last frames of the simulations and the pulling direction are shown in the rightmost column of **Figure 2**. In the following, we present the main findings regarding the stability of the 3:3 model.

#### The slow pulling reveals more fluctuation in the coiled-coil region in linear 3:3 SUN/KASH assemblies

To determine the structural fluctuation of SUN and KASH under force, we calculated the root mean square fluctuation (RMSF) of SUN2 and KASH1-5 in SUN/KASH complexes under loads **(Figure 3**). Two pulling rates (1 nm/ns & 10 nm/ns) were used to achieve the same displacement of 5 nm. Three simulations were conducted for each pulling rate and each SUN/KASH pair. The results from the three simulations were averaged for each pair. A representative image of the final frames of the simulations are shown on the right most column of **Figure 3**, displaying the final state of the SUN/KASH complexes after force application. The RMSF plots indicated that SUN2/KASH3-5 exhibit higher fluctuation in the KASH-lid region compared to SUN2/KASH1 and SUN2/KASH2. This is likely due to the proximity of the KASH-lids of SUN2/KASH3, SUN2/KASH4, and SUN2/KASH5 to the KASH domain where the loads are applied. However, both SUN2/KASH1 and SUN2/KASH2 have KASH domains that are further away from the KASH-lid. We also observed less fluctuation in the KASH domain of SUN2/KASH1 and SUN2/KASH2 compared to SUN2/KASH3-5. For all structures, the 1 nm/ns pulling rate shows more fluctuations in the coiled-coil (CC) domain than the 10 nm/ns suggesting that forces are more likely to transfer to the CC domain in the slow pulling. Moreover, the residues of CC regions for the 10 nm/ns pulling rate show a steady decrease in RMSF while the fluctuation difference over those residues is smaller for the 1 nm/ns rate (**Figure 3**). This may suggest that during the fast pulling, the forces on KASH are rapidly transferred along the protein to the end residue of the CC domain, without significant changes in the structure of the protein fragment included in our simulations. However, all CC residues experience an even distribution of stress during the slow pulling. The 3:3 versions of SUN1/KASH4 and SUN1/KASH5, obtained by splitting their 6:6 structures, were also investigated (**Supplemental Figure 1**). The RMSF distribution of the 3:3 form of SUN1 in complex with KASH4 was similar to SUN2/KASH1 and SUN2/KASH2 while the SUN1/KASH5 RMSF resembles the other three structures. The reason for this stems from KASH4 in the split version being longer (23 residues) than KASH4 (17 residues) in the SUN2 original structures. Thus, the KASH-lid region is further away from the end KASH residue.

**Figure 3:**
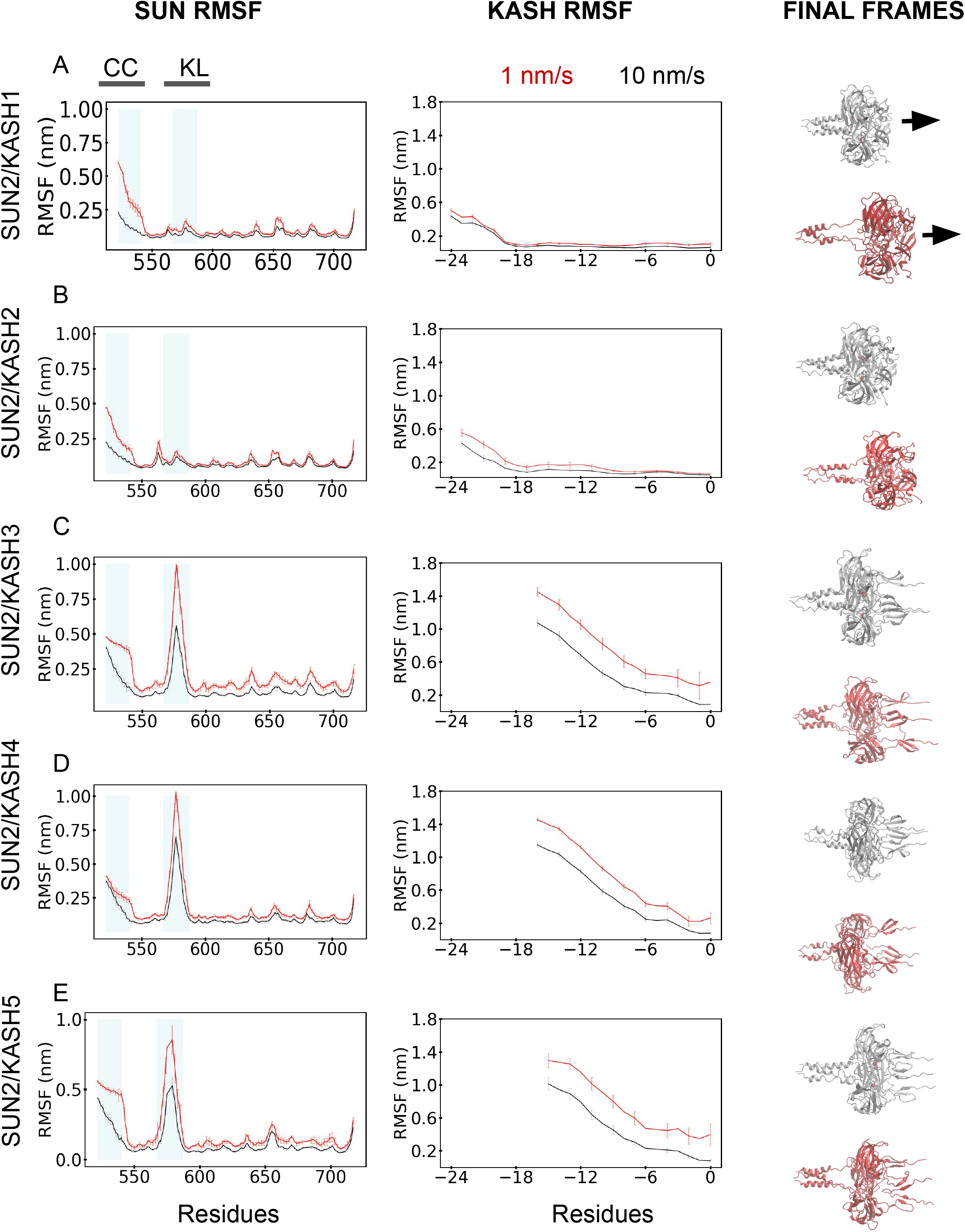
Rate dependent force response of SUN2 in complex with various KASH. Root Mean Square Fluctuation (RMSF) of SUN2 for **A)** KASH1, **B)** KASH2, **C)** KASH3, **D)** KASH4, **E)** KASH5 is shown in the **SUN RMSF** (left) column. RMSF of the KASH proteins is shown in the **KASH RMSF** (middle) column. The red and black curves represent the 1 and 10 nm/ns pulling rates respectively. The x-axis for the SUN RMSF column graphs ranges from 522 to 716 according to the SUN2 domain residue numbering. The x-axis for the KASH RMSF column graph ranges from -24 to 0 following a sequence alignment based numbering of KASH proteins^31^. **CC** represents the coiled-coil domains and **KL** represents the KASH-lid. These two areas are shaded in blue on the SUN column graphs. For each graph, the data for 3 simulations were averaged. For each simulation, the data for all protomers were averaged. Both pulling rates reach the same displacement of 5 nm. The final frames of the 10 (silver) and 1 (red) nm/ns pulling rate simulations are shown for each structure in the right column.

#### SUN2/KASH1 and SUN2/KASH2 can withstand higher forces than the other SUN/KASH pairs in the 3:3 linear model

To further investigate the reasons for the differences in RMSF, we looked at the force required to pull the various complexes apart over the trajectory of the simulation (**Figure 4A**). For both simulation rates, a larger force was required to pull SUN2/KASH1 and SUN2/KASH2 compared to the other three complexes. The force at the end of the shorter simulation is around 1300 pN for KASH1 and KASH2 and ∼400 pN for the other three structures. On the other hand, the 5 ns simulation at 1 nm/ns revealed a maximum force of ∼900 pN for SUN2/KASH1 and KASH2 and ∼330 pN for SUN2/KASH3-5. The force discrepancy confirms previous findings that SUN2 in a complex with KASH1 and KASH2 can withstand higher forces than KASH3, KASH4, and KASH5 ^34^. SUN2/KASH1 and SUN2/KASH2 exhibit a linear elastic spring behavior throughout the 10 nm/ns simulation with a mostly linearly increasing force, while the force for SUN2/KASH3-5 increases at a significantly slower rate. We notice a drop in the force intensity around 4 ns for all structures during the longer simulations. The decrease in force is more pronounced for SUN2/KASH5 compared to SUN2/KASH3 and SUN2/KASH4 which are more alike. We further investigated this force drop in the following section.

**Figure 4:**
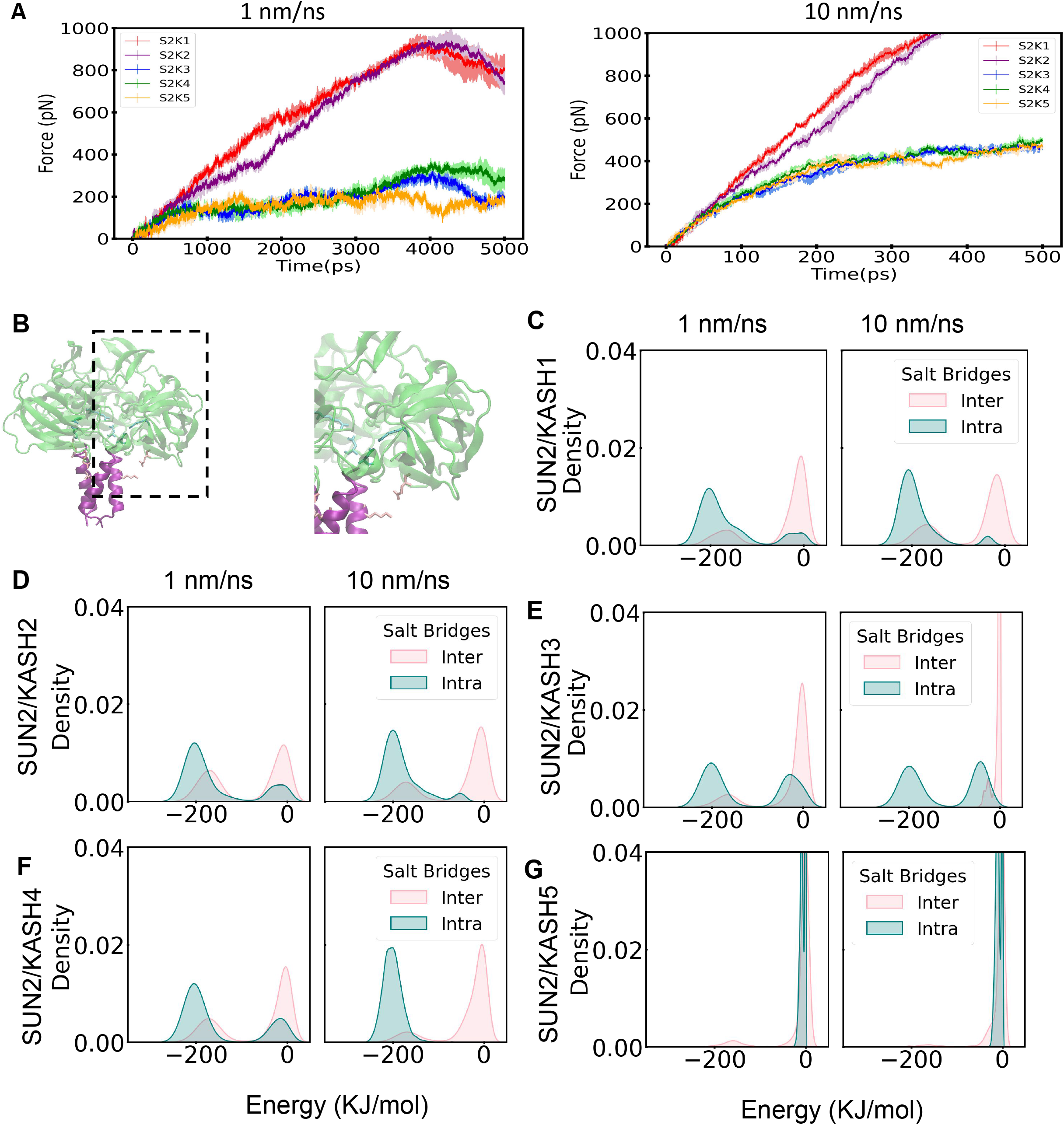
Pulling force and salt bridge interaction energies of SUN2 in complex with various KASH. **A)** Pulling force in pN over time of SUN2/KASH1 (red), SUN2/KASH2 (purple), SUN2/KASH3 (blue), SUN2/KASH4 (green), SUN2/KASH5 (brown). The left graph represents the 1 nm/ns pulling rate and the right graph shows the 10 nm/ns one. The x-axis ranges from 0 to 5000 ps for the 1 nm/ns rate and from 0 to 500 ps for the 10 nm/ns rate following the simulation time. Each simulation ran to achieve the same displacement of 5 nm. For each pulling rate and structure, three simulation data were averaged. **B)** Inter salt bridges (pink) between residues K533 of coiled-coil region and E672 of the neighboring SUN domain, and intra salt bridges (cyan) between residues D542 and R708 of each SUN protomer. The three inter and three intra salt bridges data were concatenated per type for each simulation. The y-axis shows the density and the x-axis represents the interaction energies in kJ/mol. Density plots of salt bridge interaction energies of SUN2 with **C**) KASH1, **D**) KASH2, **E**) KASH3, **F**) KASH4, **G**) KASH5. For each figure, the left graph shows the 1 nm/ns rate and the right graph represents the 10 nm/ns one.

#### Slower pulling rates result in rapid breakage of conserved intramolecular salt bridges of SUN2 in the linear SUN/KASH model

To determine the reason for a sudden drop in forces, we identified crucial residue interactions that contribute to the stability of the SUN trimer under force. We specifically looked at two important salt bridges, one linking the SUN domain of each SUN protomer to the alpha-helix (in the CC region) of the neighboring protomer (inter) and another within the SUN domain itself (intra) (**Figure 4B**). We calculated the interaction energies between residues K533 and E672 that form the intermolecular salt bridge, and between D542 and R708 that form the intramolecular salt bridge (**Supplemental Figure 2**). The salt bridge observations were concatenated per type (i.e. inter- vs. intramolecular) for each structure and simulation, and the resulting data were used to obtain density plots (**Figure 4C-G**). The main difference between the two simulation rates for all structures except SUN2/KASH5 is that all salt bridges break for the longer simulation whereas some of them remain unbroken for the shorter (**Supplemental Figure 2**). All the inter salt bridges tend to break as shown by their null interaction energies. This can better be seen on the density plots, where the density for the inter salt bridges around 0 KJ/mol is always greater than the intra salt bridges at the same energy mark (**Figure 4**). Most intra salt bridges tend to break during the longer simulations whereas they don’t break during the shorter simulations. The broken salt bridges can be seen by a higher intra density ∼0 KJ/mol for the longer simulation as compared to the shorter one. For the SUN2 KASH5 complex, most of the interactions break earlier over the course of the simulation. The short lasting salt bridges is likely due to KASH5 being the shorter KASH protomer.

### Mechanics of Force Transfer in the 6:6 SUN/KASH Complex

In the next sections, we consider the main results regarding the stability of the 6:6 or higher-order assembly of the SUN/KASH complex. The main components of the different higher-order assembly complexes are shown in **figure 5A**. We pulled on the 6:6 complex in two different directions: uniaxial and transverse (**Figure 5A, B)**. We also performed uniaxial pulling on the Apo-SUN2 structure (**Supplemental Figure 3G**). We will present the observed differences between each pulling direction, and expose our justification for these force directions by discussing some potential biological processes where SUN/KASH could possibly experience these forces.

**Figure 5:**
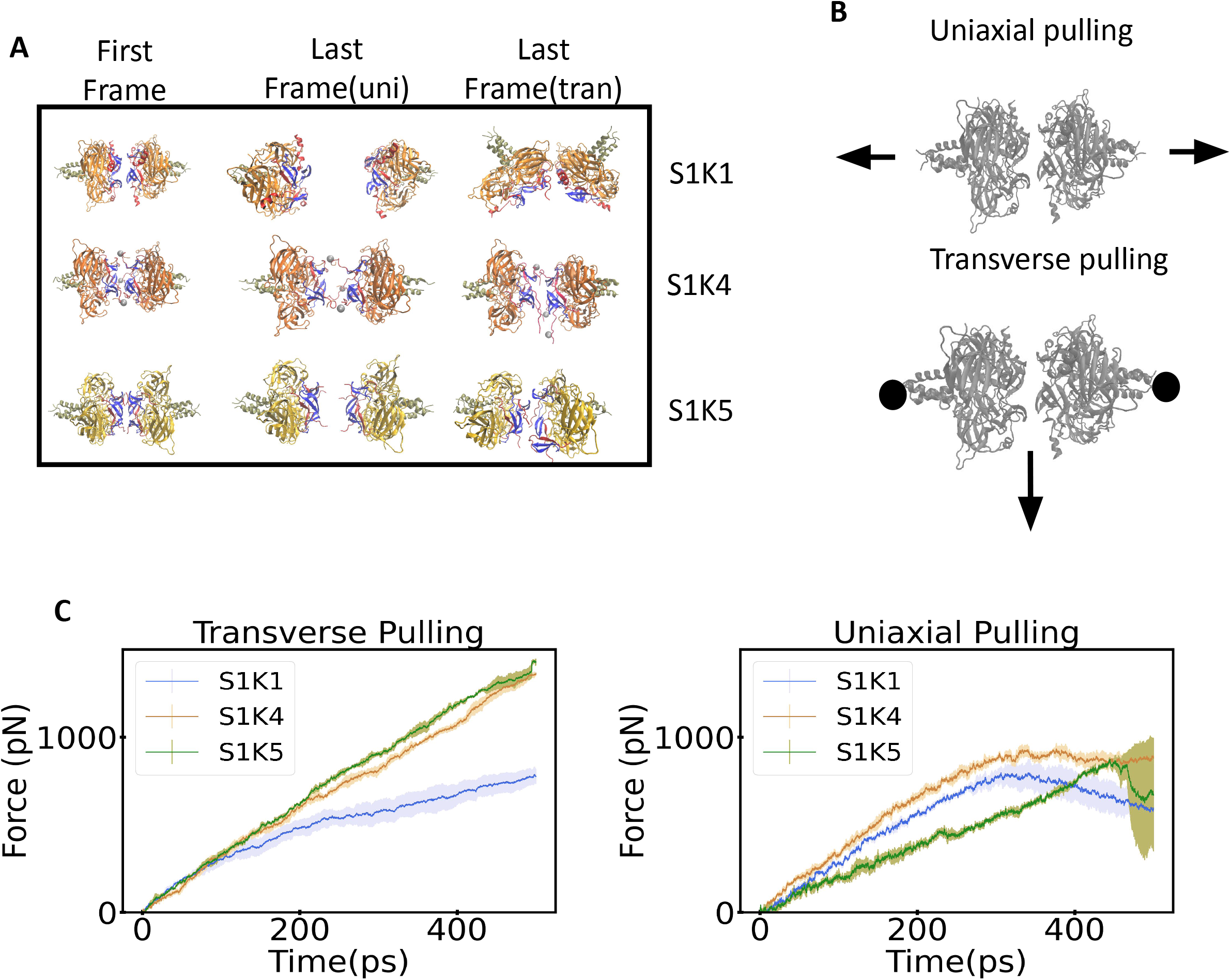
Higher-order assembly model pulling modalities and forces. **A)** First and last frame of the SUN/KASH 6:6 structures for both pulling modalities. Each is color coded to show the different regions of the protein. The blue shaded regions represent the KASH-lid within each structure. The red shaded proteins are the KASH domains. The orange shaded region is the SUN domain. And the gold shaded region are the beta sheet folds of SUN. **B)** Depiction of uniaxial and transverse pulling. **C)** A comparison of the amount of force used throughout the simulation for each pulling modality over simulation time.

#### SUN/KASH hexamers completely dissociate in SUN1/KASH1 under uniaxial pulling (but not in SUN1/KASH4 and SUN1/KASH5)

Contrasting the initial and final frames of the molecular dynamics trajectory of the uniaxial pulling simulation, we can infer that the separation of KASH heads for SUN1/KASH1 is more appreciable than for SUN1/KASH4 and SUN1/KASH5 (**Figure 5C**). Pulling the SUN/KASH heads apart requires the same force magnitude until 0.3 ns. At 0.3 ns of the simulation, the SUN1/KASH1 trimers completely separate, while SUN1/KASH4 and SUN1/KASH5 still maintain some of their head-to-head interactions. The lack of complete dissociation is shown in the middle column of **Figure 5A**, where there are clear differences between the distances separating KASH heads. The structural differences between the SUN/KASH systems (**Figure 5A)**, are responsible for discrepancies in separation. The same could be said for slower pulling rates. The amount of force required to break the head-to-head interaction is smaller, but the overall trends remain the same. The next pulling modality we looked at is transverse pulling.

#### SUN1/KASH1 experiences a more distinct conformational change than SUN1/KASH4 and SUN1/KASH5 but less force under transverse pulling

In the transverse pulling modality, we applied forces to the end of each KASH instead of SUN. The transverse pulling modality showed some distinct conformational changes within the different SUN/KASH structures (see **Figure 6A)**. The force required to pull on KASH was noticeably greater in SUN1/KASH4 and SUN1/KASH5 as compared to SUN1/KASH1 (**Figure 5D)**. The SUN1/KASH1 structure reaches a maximum load of 708 pN under transverse pulling while SUN1/KASH4 and SU N1/KASH5 attain a maximum force of 933 pN and 1365 pN, respectively, for transverse pulling (**Figure 5D)**. The differences in maximum force between SUN1/KASH1 and SUN1/KASH4,5 are directly related to the head-to-head interaction. However, before we examine the head-to-head interaction between the different structures, we should quantify the structural changes experienced by each SUN/KASH complex. The change in angle between CC regions on opposing SUN trimers was used to calculate the overall angle change of the structure. A simplified version of all 6:6 structures in the first and last frames of transverse pulling and the rate of angle change over the simulation time are presented in **Figure 6A, B**. SUN1/KASH1 shows the greatest angle change while, interestingly, SUN1/KASH4,5 show relatively the same angle change over the simulation time. One may expect the SUN/KASH system to start at roughly 180°; however, in their original structures, the opposing CC regions are not coaxial. SUN1/KASH1 changes 40° over the simulation time. On the other hand, SUN1/KASH4,5 changes ∼5° over the simulation time. Like the other results regarding the 6:6 structure, the angle change can be explained by the differences in head-to-head interaction of the three structures.

**Figure 6:**
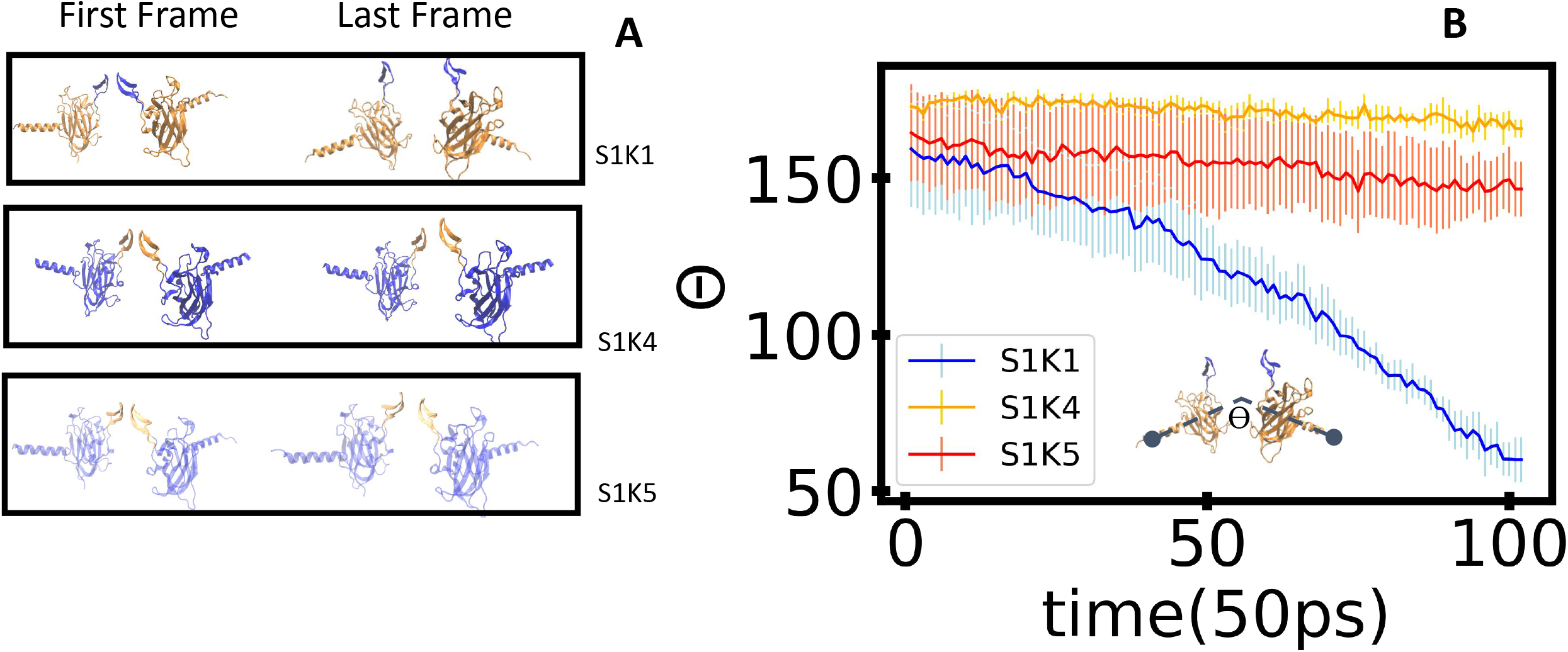
Conformational changes within SUN/KASH structures under transverse pulling. **A)** The first and last frames of transverse pulling for all 6:6 structures. Only two protomers were used to show the change in conformation. **B)** Angle changes between alpha helices of adjacent SUN protomers over the simulation time (Ө) as shown in the inset schematic.

To determine whether the head-to-head interactions play a role in maintaining the integrity of the structures, we used both GROMACS interaction energies and a pairwise distance heatmap (**Figure 7**). We focused on 2 residues, 671 and 673, on the KASH-lid because these residues are thought to primarily maintain head-to-head KASH-lid interaction. For SUN1/KASH1, there was a steady drop in interaction energies between KASH-lids under force (Figure 7A). The sudden drop of energy corresponds to the same pairwise distance between residue 671 of corresponding KASH-lids (Figure 7D). In SUN1/KASH4, the KASH-lid does not participate in bonding; this can be seen on the heatmap (**Figure 7E)**, and the near zero interaction energies shown in **Figure 7B**. In the case of SUN1/KASH4, the main interactions that hold the structure together are the zinc coordination bonds with KASH. Finally, **Figure 7C** shows the different interactions between the various KASH-lid pairs in SUN1/KASH5. In SUN1/KASH5, a breakage of the interaction between residues 545, which corresponds to the PPP motifs is observed as evident from the heatmap in **Figure 7F**.

**Figure 7:**
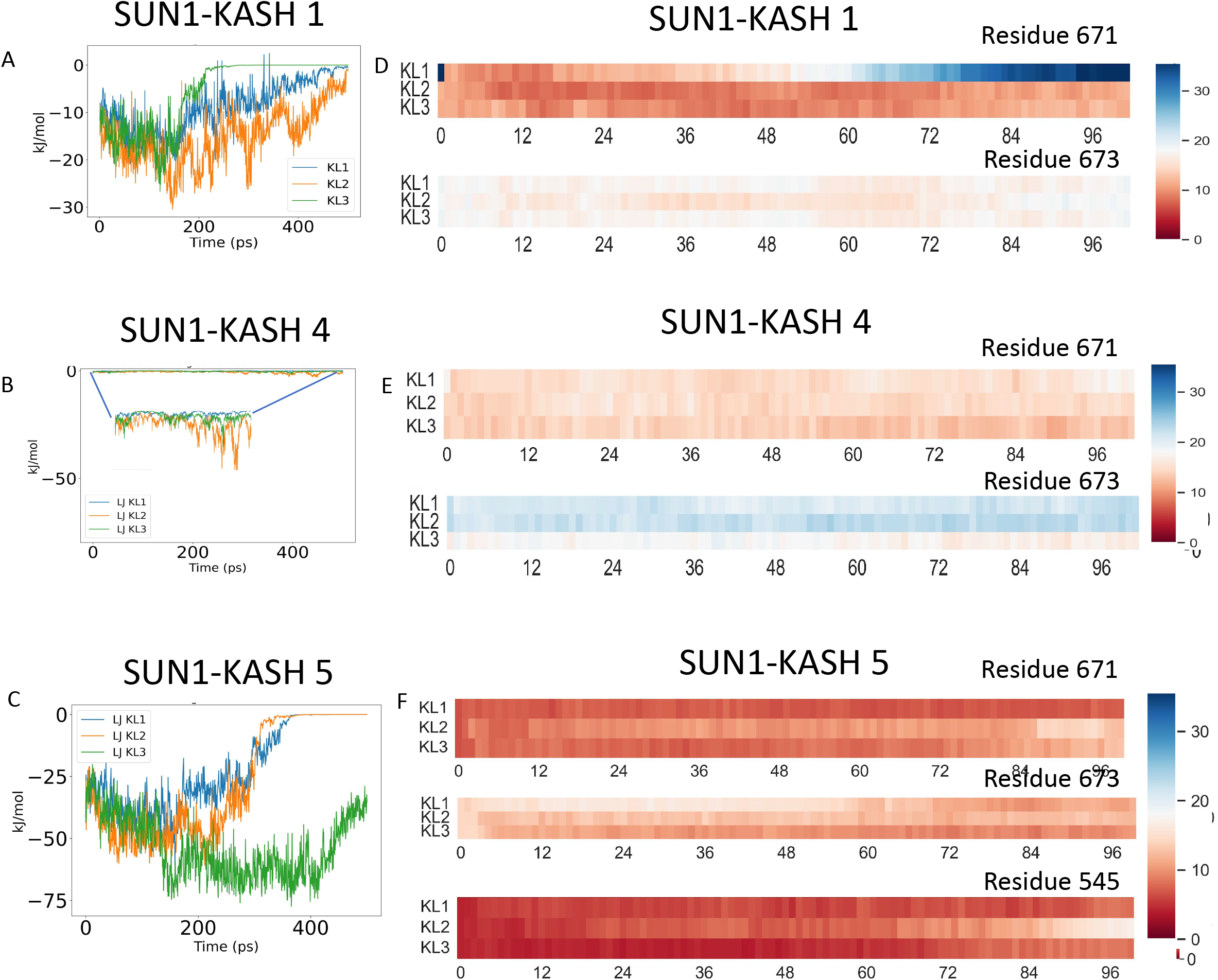
Interaction energies between KASH-lids of various SUN/KASH pairs. **A)** The Lenard Jones (LJ) interaction energies between KASH-lid (KL) pairs (i.e. KASH-lid of a protomer on one SUN trimer with adjacent KASH-lid of a protomer on other SUN trimer) for **A)** SUN1/KASH1 **B)** SUN1/KASH4, and **C)** SUN1/KASH5. **D)** Heatmaps of the pairwise distances between the alpha carbons of residues 671 or 673 of adjacent KASH-lids on opposite SUN trimers over the simulation time for SUN1/KASH1 and **E)** SUN1/KASH4. **F)** Heatmaps of the pairwise distances between the alpha carbons of residues 671, 673 and 545 of adjacent KASH-lids on opposite SUN trimers over the simulation time for SUN1/KASH5.

#### Apo-SUN2 main head-to-head interaction requires considerably less force to break

As control simulation, the force response and interactions within the Apo-SUN2 structure was examined next. Apo-SUN2 refers to the trimer of SUN2 containing a KASH-lid region but no KASH associated. Since both SUN1 and SUN2 have very similar structures, we assumed that the forces experienced and the residues that contribute to the head-to-head interactions are similar. We first decided to compare the amount of force required to split the structure apart (**Supplemental Figure 3)**. The Apo structure requires considerably less force to break the head-to-head interaction than the SUN/KASH complexes. This result is consistent across the different pulling rates. Additionally, we considered the interaction energies between the KASH-lids of the different Apo heads. We measured the Lennard Jones short-range interaction energies between the opposing KASH-lids, given the important role they play in SUN1/KASH1 head-to-head interaction. Our results suggest the KASH-lid also plays a significant role in maintaining the Apo head-to-head interaction (**Supplemental Figure 3**).

### Force and RMSF Comparison of the 3:3 and the 6:6 models

The force response and the RMSF were used to compare the uniaxial pulling modality of the linear trimer model and the transverse pulling modality of the higher-order assembly. We believe comparing both models is appropriate, since we are effectively pulling on the KASH protomers in these two modalities. The force over the simulation time between the two models differs greatly. The SUN/KASH complexes in the 3:3 form that withstand higher forces withstand lower forces in the 6:6 model. We compared the structures in the linear trimer model to their counterparts in the higher-order assembly based on the KASH, because of the similarity between SUN1 and SUN2. Thus, SUN2/KASH1 (3:3 model) experiences a lot more force compared to the 6:6 assembly of SUN1/KASH1 for the same displacement. On the other hand, SUN2/KASH4,5 experience less force in the linear trimer form than the 6:6 model of SUN1/KASH4,5. Both of these phenomena can be explained through the structure of the different SUN/KASH variations and pulling directions.

We also compared the RMSF between the different models. We aligned SUN1 residue numbering to SUN2, following other previous studies on comparing the similarities between SUN1 and SUN2 ^43^. As seen in **Figure 8**, the CC region, which is between 522-540, experiences a greater fluctuation in the 3:3 model than in the 6:6 one. The other noticeable change occurs in the KASH-lid region. SUN2/KASH3-5 show large RMSF values in residues 567-587 (KASH-lid region in SUN2), while SUN2/KASH1,2 and SUN1/KASH1,4,5 do not exhibit appreciable structural fluctuations. The CC region is closely connected, in terms of residue spacing to the KASH-lid region. Also, by pulling on KASH we are effectively pulling on the KASH-lid. Consequently, the CC region affects the KASH-lid and vice versa. Overall, SUN1/KASH1,4,5 and SUN2/KASH1,2 do not possess high RMSF compared to SUN2/KASH3-5. In the case of SUN2/KASH1,2, this difference can be explained by the presence of disulfide bonds which prevent high fluctuations in KASH-lid and CC regions, thereby maintaining the structural integrity of the complexes under high load. For the SUN1/KASH1,4,5, other interactions are keeping the SUN/KASH higher-order assembly relatively stable as KASH is pulled in a different direction.

**Figure 8:**
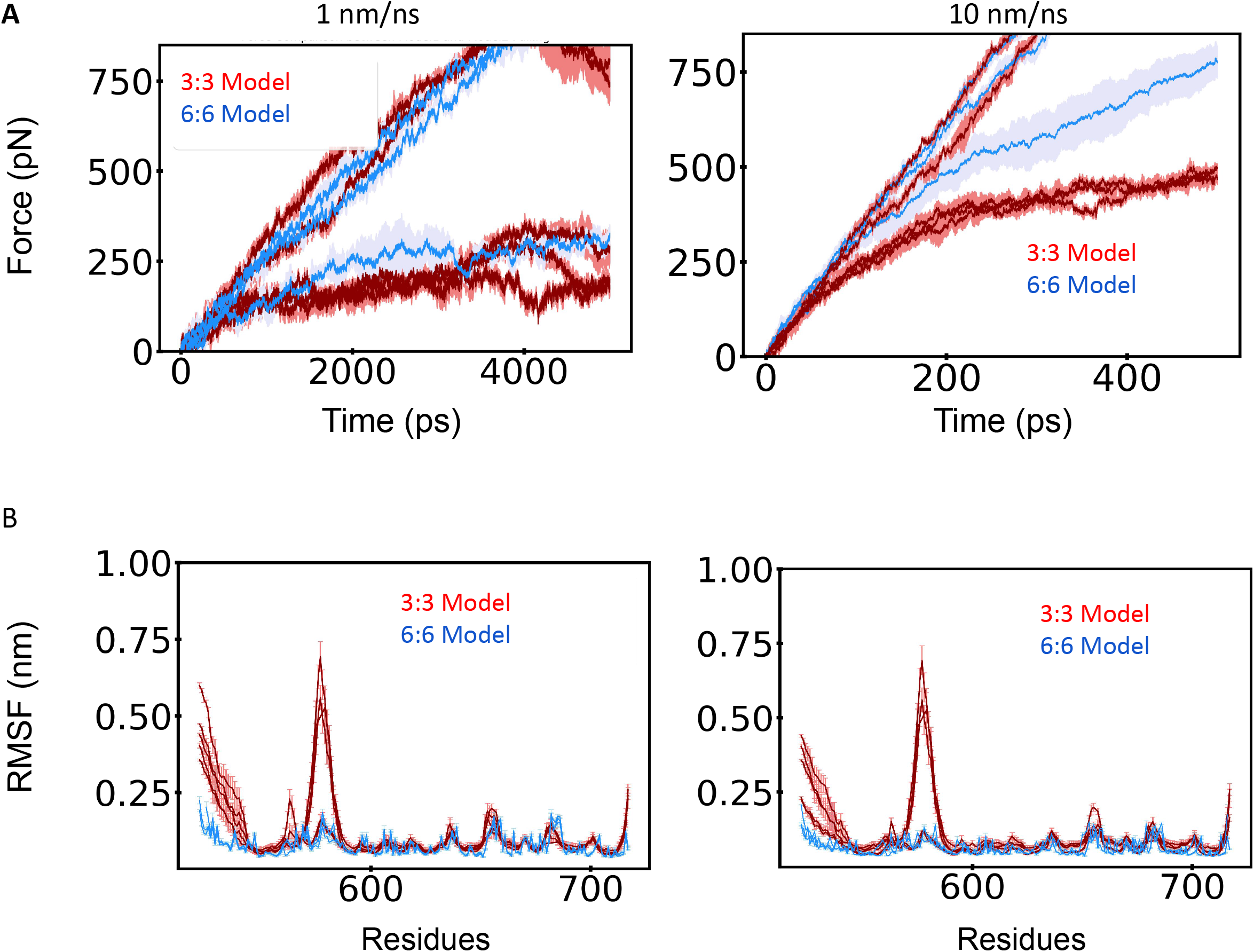
Force and RMSF comparison between the two different models of SUN/KASH over the simulation time. The mode of pulling we are using to compare for the 6:6 structure is transverse pulling. **A)** The 6:6 structures are all colored blue while the 3:3 structures are colored red. The standard error is shaded in a lighter hue. Unlike 3:3 the standard error in 6:6 noticeably increases. With the exception of SUN2/KASH1,2, the 6:6 structures (blue) experience more force than the 3:3 structures. The differences between 3:3 and 6:6 are the increased size of the standard error and the noticeable decrease in force in the latter half of the simulation. The forces in 3:3 all tend to decrease during the end of the simulation and the forces in 6:6 tend to increase over the simulation time. **B)** All of the 6:6 structures are colored in blue while all of the 3:3 structures are colored in red. The residues for 6:6 (contained SUN1) were shifted to match residue numbering in 3:3 (SUN2). The main differences between the two arrangements are the CC region and the KASH-lid region. 3:3 structures have a noticeable peak for SUN2/KASH3,4,5, and a relatively lower peak for all other structures. 3:3 has higher fluctuations in the CC region compared to 6:6.

## Discussion

Recent higher-order assembly models for the LINC complex have debated the putative mechanism of force transfer within the SUN/KASH models. Using molecular dynamics simulations we set out to compare and contrast the putative 3:3 SUN/KASH complex model vs. the new higher-order assembly 6:6 SUN/KASH complex. Here we discuss the implications of our findings for each model in terms of their structural biomechanics along with their contribution to a broader functional level. We argue that both models can assemble in unique networks and may exist simultaneously in the cell depending on the tasks performed.

Besides anchoring the nucleus, the LINC complex serves as a load bearer and a force transmission agent ^34^. Its CC region provides the ability to behave elastically as a spring under force. However, the rate at which these forces are applied plays a significant role in the force transmission mechanism and thereby the stability of the structure. Our results suggest the existence of SUN/KASH clustering in the nucleus for both models.

We showed that SUN/KASH in the 3:3 arrangement is force rate dependent. Forces are either concentrated and transmitted to the opposite end of the complex or transmitted throughout the entire SUN domain. We also demonstrated the previous postulates on SUN2/KASH1,2 being able to withstand higher forces than other 3:3 structures. Other parameters may play a role in the differences observed in the force transmission mechanism of slow and fast pulling rate. We hypothesized that salt bridges between CC and SUN domain as well as within the SUN protomers contribute chiefly to the force mechanism. Our results indicate that they tend to break more easily when forces are applied at a slower rate, leading to the higher fluctuations observed in the structures. When the forces are applied over a shorter time period, these salt bridges do not experience stress immediately and some remain intact for the same displacement. The recruitment of SUN/KASH to an area of the nucleus may still be a mystery, but the cleavage of salt bridges and increased fluctuation suggests that under a network of SUN/KASH the fluctuation will decrease. A SUN/KASH cluster would also be force rate dependent like a single unit of SUN/KASH.

Like the 3:3 arrangement, the 6:6 arrangement results show that different SUN/KASH complexes respond differently to force. The biggest differences in the higher-order assembly by far are between SUN1/KASH1 and SUN1/KASH4,5. A rationale of this difference pertains to the disulfide bonds between SUN1 and KASH1. Regardless of the model discussed, the disulfide bonds remain an integral part of the structures that we have studied ^31,34,44^. SUN1/KASH1 also seems like the most probable SUN/KASH system to exist in the nuclear envelope in both 3:3 and 6:6 arrangements. A legitimate concern can be raised about the way SUN1/KASH1 in the 6:6 arrangement handles force if the head-to-head interaction does not hold at larger forces. KASH1 and KASH2 are meant to interact with actin and other cytoskeletal elements ^15,16^. Thus, it is reasonable to think that the SUN/KASH will always experience some tension. Under relatively small loads, SUN1/KASH1 may be in the 6:6 arrangement and when the cell experiences more loads over time, it may be in the linear trimer form. These speculations can be cleared up if we had a better understanding of the CC region. In other higher-order SUN/KASH structures, it is shown that SUN1 could potentially assemble adjacent to each other instead of the head-to-head interaction that we have shown ^43,45,46^. This conformational change could potentially happen with SUN1/KASH1 under extreme cellular events such as mitosis or apoptosis ^47,48^.

The two other 6:6 structures, namely SUN1/KASH4,5, can experience more force without undergoing as much conformational change as SUN1/KASH1. KASH5 is known to be responsible for meiotic processes, specifically chromosomal movement ^49^. While KASH5 is the smallest KASH protein we have observed, it experiences more force under pulling in the 6:6 arrangement. The lack of conformational change leads us to assume how it orientates itself in the nucleus in higher-order networks or SUN/KASH clusters. KASH4 is usually expressed in secretory epithelial cells ^16^. Many studies have shown the importance of zinc in epithelial cells ^50^. KASH4 also exclusively binds to kinesin-1, which plays a role in nuclear positioning. The 6:6 KASH4 has zinc as part of its structure. There may be some mediated action that potentially releases zinc from SUN/KASH, in order to perform specific tasks.

As there is not as much of a conformational change seen in the SUN1/KASH4,5, one may wonder how these structures localize in the nuclear membrane, and what network SUN and KASH make in the nuclear envelope. Even though we have the crystal structure of SUN/KASH, we do not have a clear understanding of the CC region that spans the perinuclear space. Since the 6:6 KASH4 and KASH5 do not undergo any drastic conformational changes, it would be unrealistic for the CC region to bend 90 degrees without other protein elements. Different arrangements of SUN/KASH can be postulated in the nucleus as depicted in **Figure 9**. These SUN/KASH arrangements cluster to form unique symmetries that can span the nuclear envelope. Both 3:3 and 6:6 arrangements cluster in various configurations (**Figure 9B**,**C)**. As previously stated, we recognize that both 3:3 and 6:6 arrangements can exist simultaneously in the nucleus, therefore a hybrid version of both arrangements is also conceivable in the nuclear envelope (**Figure 9D**). Thus, it would be likely that there exist elements along the SUN coiled-coil region that add to the mechanics of SUN/KASH and contribute to a greater meshwork of the nuclear envelope.

**Figure 9:**
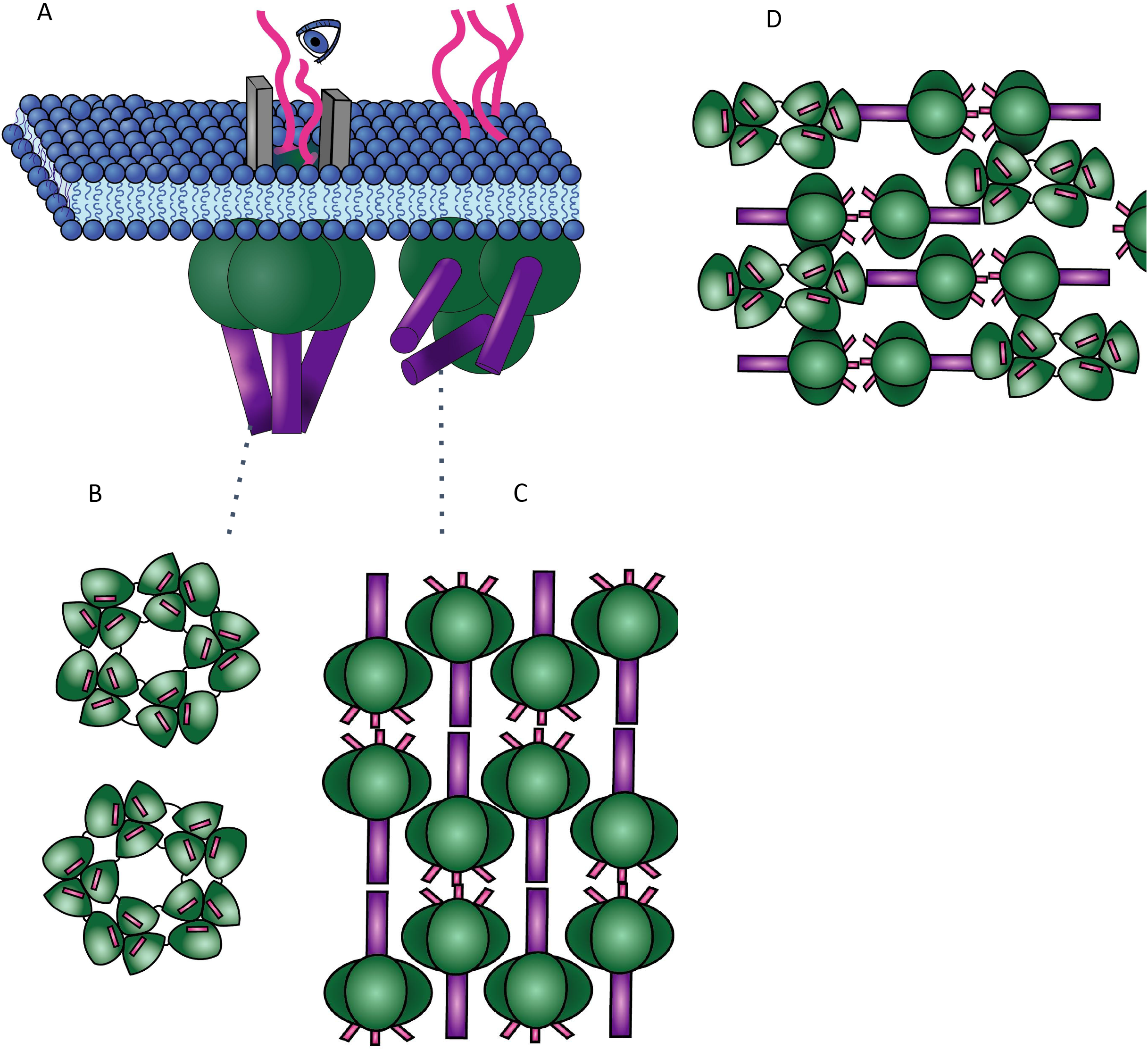
Proposed network of higher-order clustering of different SUN/KASH complexes. **A)** An orthographic cartoon depicting the potential ways both 3:3 structures and 6:6 structures may associate in the outer nuclear envelope. The KASH (pink) extends from the outer nuclear envelope (blue) while SUN sits on the underside. **B)** A top view depiction of how the 3:3 SUN/KASH complexes form a unique star shaped network. **C)** A representation of a top view of 6:6 SUN/KASH structure interacting to form a matrix network. **D)** A representation of how both 3:3 and 6:6 structures come together to form a uniquely packed matrix network.

The force required to pull apart Apo-SUN2 is considerably smaller than all other SUN/KASH complexes. This may functionally suggest a few things. First, Apo-SUN2 may have a special recruitment pathway where, both maintaining the head-to-head interaction and detaching from the head-to-head is important for switching between the two arrangements. The ability to attach and detach also could play a role in nuclear membrane localization. Finally, it may imply how important KASH is for distributing and withstanding higher forces.

We have shown the importance of different pulling rates and directions in the mechanobiology of the LINC complex. The central question of this study was to determine what SUN/KASH arrangement is more likely to exist in the cell and how these complexes may cluster in the nucleus. We believe that both arrangements may exist simultaneously depending on the type of cells, the stage of the cell cycle, and the functions performed. The linear trimer model may be suitable for fast tensile force transmission and high loads, while the higher-order assembly model may be suitable for describing the interaction between SUN and KASH under compressive and shear forces. We also suggest that when these clusters form, they augment and amplify the structural rigidity and dynamics of the nuclear envelope. Nonetheless, we do not have a clear understanding of the other components that may be involved in the force transfer mechanisms, especially in the case of SUN1/KASH1. These binding partners of the LINC complex should be investigated to better grasp the extent of their influence on the arrangement of the SUN/KASH complexes.

## List of Abbreviations

LINC: LInkers of Nucleoskeleton and Cytoskeleton
CC: Coiled coil
S2K1/2/3/4/5: SUN2 protomer in complex with KASH1/KASH2/KASH3/KASH4/KASH5
S1K1/4/5: SUN1 protomer in complex with KASH1/KASH4/KASH5
Apo-SUN2: Trimer of SUN2 with no KASH associated
NE: Nuclear Envelope

## Figure captions

**Supplemental Figure 1:**
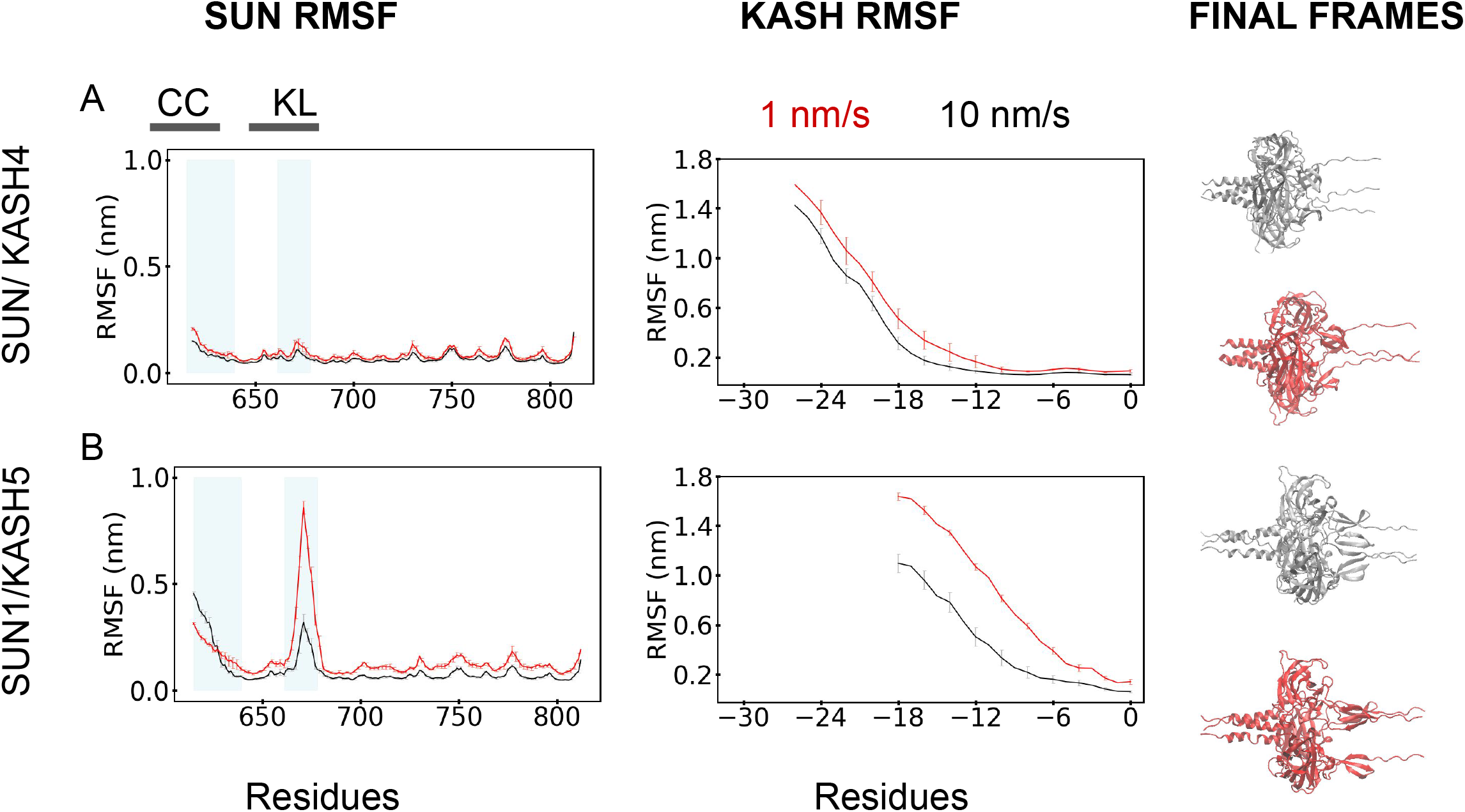
Rate dependent force response of SUN1 in complex with KASH4,5. Root Mean Square Fluctuation (RMSF) of SUN1 for **A)** KASH4, **B)** KASH5 are shown in the SUN RMSF (left) column. RMSF of the KASH proteins is shown in the KASH RMSF (middle) column. The red and black curves represent the 1 and 10 nm/ns pulling rates respectively. The x-axis for the SUN RMSF column graphs ranges from 522 to 716 according to the SUN2 domain residue numbering. The x-axis for the KASH RMSF column graph ranges from -24 to 0 following a sequence alignment based numbering of KASH proteins ^31^. **CC** represents the coiled-coil domains and **KL** represents the KASH-lids. These two areas are shaded in blue on the SUN column graphs. For each graph, the data for 3 simulations were averaged. For each simulation, the data for all protomers were averaged. Both pulling rates reach the same displacement of 5 nm.The final frames of the 10 (silver) and 1 (red) nm/ns pulling rate simulations are shown for each structure in the right column.

**Supplemental Figure 2:**
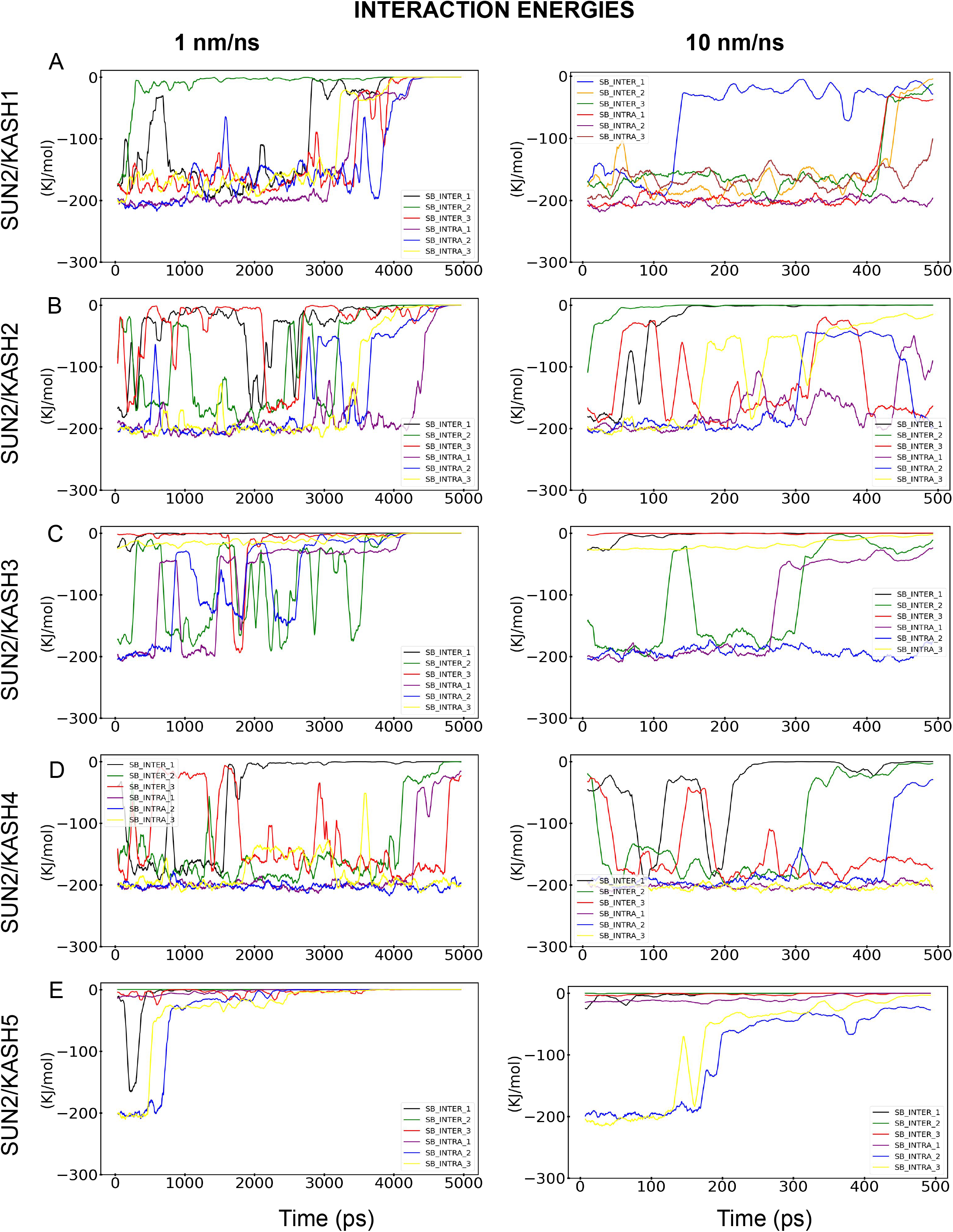
Interaction energies between salt bridges in SUN2 structures. Short-range coulombic interaction energies of SUN2 in complex with **A)** KASH1, **B)** KASH2, **C)** KASH3, **D)** KASH4, **E)** KASH5 are shown for the 1 nm/ns rate (left column) and 10 nm/ns (right column). SB-inter 1-3 denotes the three intermolecular salt bridge pairs in each structure and the SB-intra 1-3 shows the three intramolecular salt bridges. The x-axis displays the simulation time in ps, while the y-axis shows the energy in kJ/mol. For each structure and pulling rate, only one sample simulation energy is shown.

**Supplemental Figure 3:**
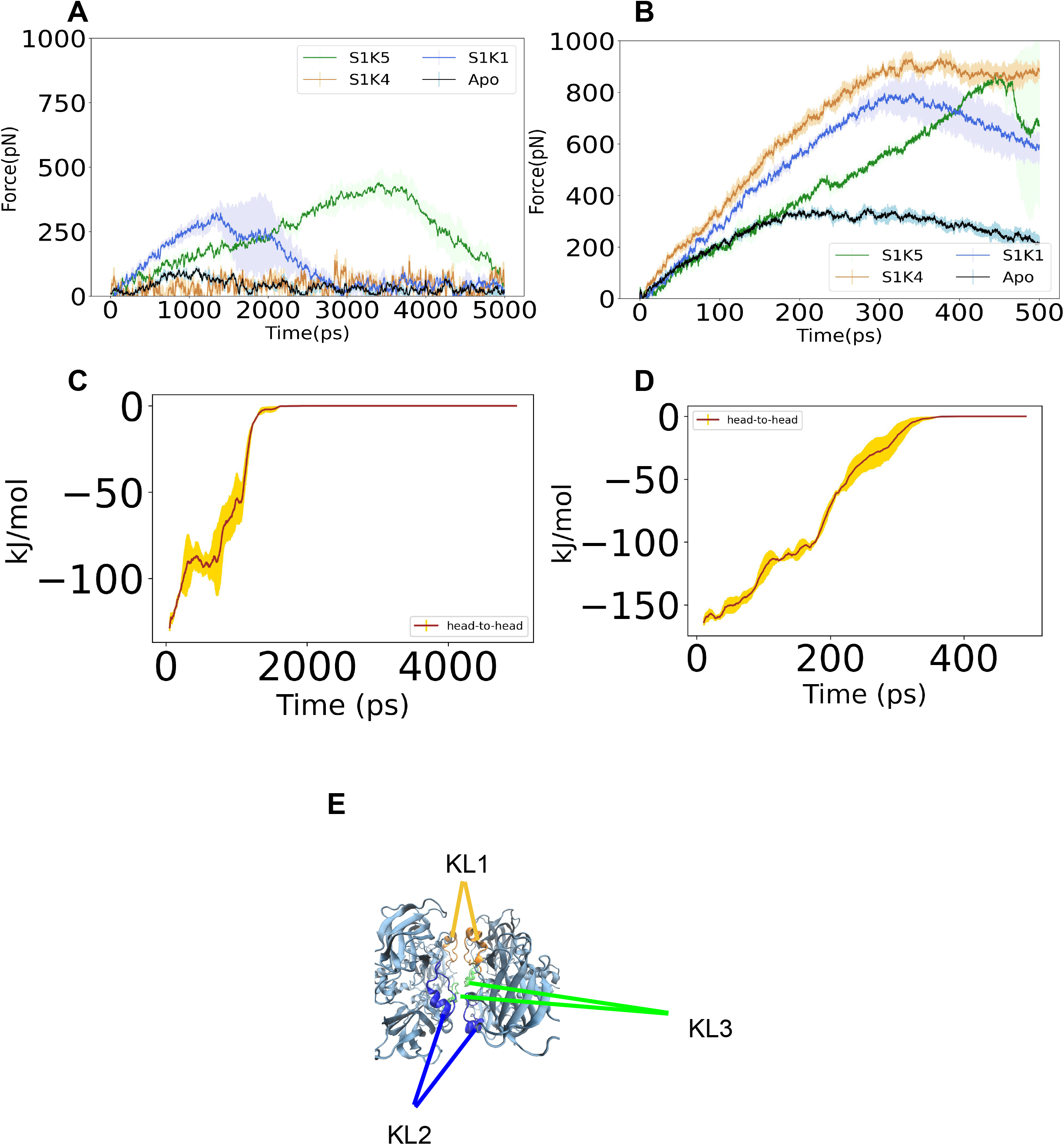
Comparison of Apo-SUN2 to other 6:6 structures. Both **A)** and **B)** show the uniaxial pulling force in different pulling rates. Apo-SUN2 structure, under slow pulling, experiences less force before it dissociates. The structures respond to different pulling rates. **C)** and **D)** are looking at the head-to-head interaction energy for different pulling rates over three simulations. The total sum of the KASH-lid pairs equals the total amount of energy seen in **C)** and **D). E)** is a diagram showing the KASH-lids responsible for head-to-head interaction.

**Supplementary Figure 4:**
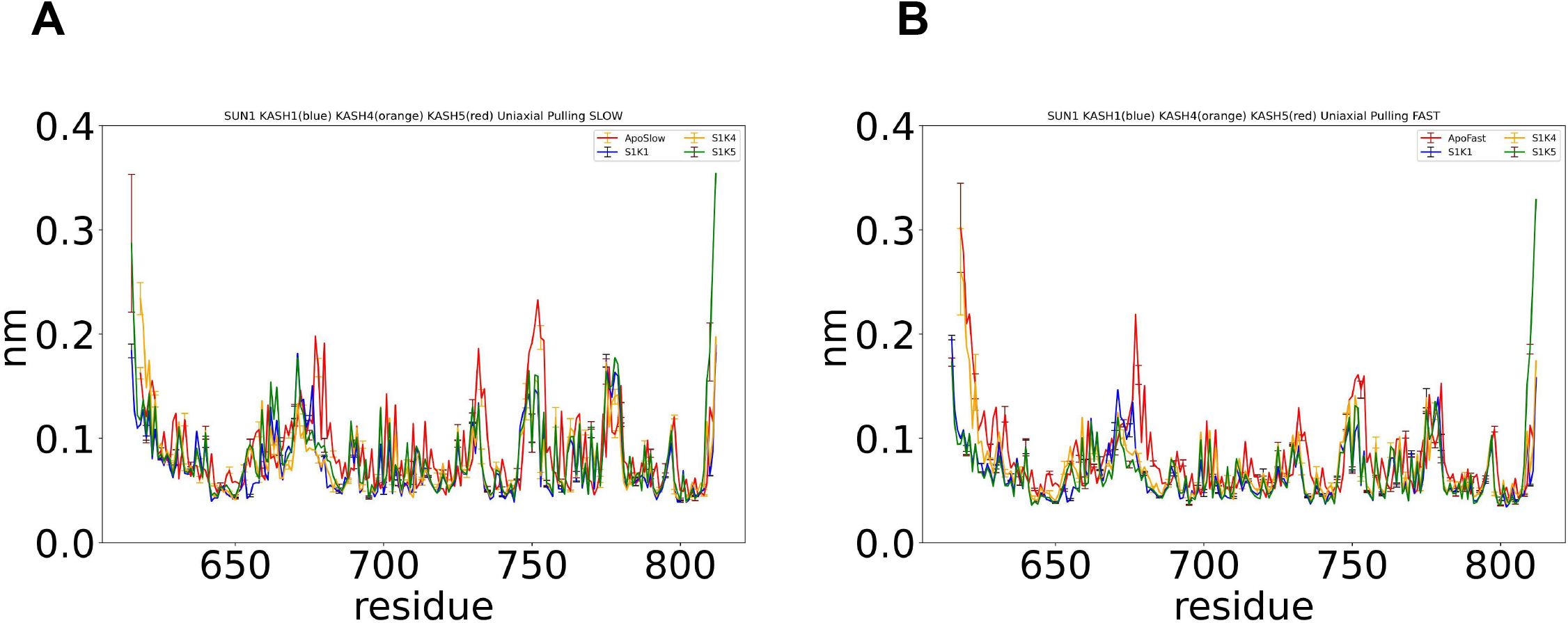
Comparing the RMSF between APO and other 6:6 structures for **A)** slow pulling rate and **B)** fast pulling rate

